# A MYB transcription factor, *PlMYB308*, plays an essential role in flower senescence of herbaceous peony

**DOI:** 10.1101/2021.09.02.458764

**Authors:** Xiaotong Ji, Zhuangzhuang Xu, Meiling Wang, Xuyang Zhong, Lingling Zeng, Daoyang Sun, Lixin Niu

## Abstract

Herbaceous peony is an important cut-flower plant cultivated across the world, but its short vase life substantially restricts the economic value of this crop. It is well established that endogenous hormones regulate the senescing process, but the molecular mechanism of them in flower senescence is still unclear. Here, we isolated a MYB transcription factor gene *PlMYB308* from herbaceous peony flowers. Transcript abundance of *PlMYB308* was strongly up-regulated in senescing petals. Silencing of *PlMYB308* resulted in delayed peony flower senescence, and dramatically increased gibberellin (GA) but reduced ethylene and abscisic acid (ABA) levels in petals. Ectopic overexpression of *PlMYB308* in tobacco accelerated flower senescence, and reduced GA but increased ethylene and ABA accumulation. Correspondingly, biosynthetic genes of ethylene, ABA, and GA showed variable expression levels in petals after silencing or overexpression of *PlMYB308*. A dual-luciferase assay showed that PlMYB308 specifically bound to the *PlACO1* promoter. High expression levels of *PlMYB308* were accompanied by low petal anthocyanin accumulation in senescing petals. A further bimolecular fluorescence complementation assay revealed an interaction between PlMYB308 and PlbHLH33, which was supposed to inhibit the anthocyanin biosynthesis. Taken together, our results suggest that the PlMYB308-*PlACO1* and PlMYB308-PlbHLH33 regulatory checkpoints perhaps positively and negatively operate the production of ethylene and anthocyanin, respectively, and thus contribute to the senescence with impaired pigmentation in herbaceous peony flowers.

## Introduction

Senescence is a complex coordinated process in the whole organ, tissue, and cellular levels (Tripathi et al., 2014). In agricultural production, particularly in transportation and storage, senescence dramatically decreases crop yield and causes economic loss in commercial crops and ornamental plants (Gan and Amasino, 1997). Nowadays, people have used physiological, biochemical, and molecular approaches to study the senescence process in various plant organs, such as fruits (Kou et al., 2012), leaves (Zhang and Gan, 2012), and flowers (Chen et al., 2011). The phenotypic changes of senescence such as color fading or wilting are mediated by complex pathways, and the degradation of polymer or nucleic acids and proteins biosynthesis can influence these pathways. It has been reported that many up- or down-regulated structural genes and transcription factors (TFs) are implicated in the plant senescence (Guo and Gan, 2012).

Abiotic and biotic stresses may trigger flower senescence. Numerous studies have provided evidence that there is an intricate and complex network among phytohormones (Zhang and Zhou, 2013). It has been well known that ethylene is an important endogenous hormone which regulates the flower senescence. A burst of ethylene production can cause flower senescence in many plants such as petunia and carnation (Woodson et al., 1992; Tang et al., 1994). Treating these plants with exogenous ethylene accelerates their flower senescence while the treatment with ethylene biosynthesis inhibitors retards the process (Hark, Dekker, and Essers, 1991; Serek, Sisler, and Reid, 1995). It has been revealed that in daylily, ABA is a primary hormone that regulates the flower senescence, but this process may be stimulated by ethylene accumulation (Panavas et al., 1998). By contrast, GA or cytokinin can delay flower senescence (Saks et al., 1992). Some researchers have attempted to understand the senescence-modulating network and constructed enriched cDNA libraries to identify differentially-expressed transcripts at different stages of flower senescence in daylily, carnation, and daffodil (Lawton et al., 1989; Valpuesta et al., 1995; Hunter et al., 2002). However, the cross-talk among plant hormones and the complex signaling pathways during flower senescence are still unclear.

The senescence behavior of many floral tissues is concomitant with the petal color fading due to a decrease in anthocyanin content. Anthocyanin biosynthesis primarily involves a series of structural genes and TFs known as MYB, bHLH, and WD40 repeat proteins, which commonly form a trimeric MYB-bHLH-WD40 (MBW) complex to regulate certain downstream anthocyanin-related structural genes (Gonzalez et al., 2010). The MYB proteins comprise a large and functionally diverse TF family in all eukaryotes. They have highly conserved MYB domains binding to specific DNAs. In Arabidopsis, the MYB proteins function as critical mediators of many different biological activities, suggesting that they have extensive functional diversification. In the past few decades, many MYB TFs have been characterized to be participated in various biological processes, including plant development, cell shaping, hormone signaling, and biotic or abiotic stress tolerance. For instance, the separation of lateral organ and the formation of axillary meristem can be controlled by Arabidopsis AtMYB105 and AtMYB117 upstream of AtMYB37 (Lee et al., 2009). Transcriptional activation of *AtMYB96* contributes to the plant tolerance to drought stress by mediating ABA signaling (Seo et al., 2009) and the biosynthesis of cuticular wax in Arabidopsis (Seo et al., 2011). ABA treatment induces the expression of *AtMYB2* (Shinozaki et al. 1992). It has also been reported that *AtMYB30* controls hypocotyl cell elongation in seedlings by participating in the brassinosteroid pathway (Li et al. 2010). The R2R3 MYB TF is an important component in the MBW complex which affects anthocyanin accumulation (Stracke et al. 2001). Although there are many observations about MYB proteins, the function of MYB308 TFs in hormone-regulated petal senescence and anthocyanin biosynthesis of herbaceous peony remains unknown.

The economic value of herbaceous peony cut flowers largely depends on excellent flowering quality and long vase life. Thus, the regulatory mechanism of flower senescence is one of the important topics in postharvest physiological research of herbaceous peony. In this study, we reported that *PlMYB308*, a member of MYB family, participated in the accumulation of ethylene and anthocyanin in herbaceous peony petals. Its silencing in herbaceous peony and overexpression in tobacco increased and decreased the flower longevity, respectively, supporting a crucial role of PlMYB308 in the regulation of flower senescence.

## Results

### Expression of *PlMYB308* increases during flower senescence

The full-length cDNA sequence of *PlMYB308*, containing a 759-bp open reading frame (ORF) region, was isolated from the floral tissues of herbaceous peony. Through the BLAST search against non-redundant GenBank databases in the NCBI, the highly homologous proteins to the deduced polypeptides encoded by *PlMYB308* gene were obtained. Amino acid alignment and phylogenetic analysis showed that PlMYB308 was relatively close to the MYB308s from other plant species, such as *Salvia miltiorrhiza* MYB308 (AGG09847.1), *Vitis vinifera* MYB308 (NP_001268129.1), *Rosa hybrid* MYB308 (XP_022724681.1), and *Petunia hybrida* MYB308 (ADX33331.1) (Figure 1A). All the homologous sequences exhibited high conservation of amino acids in the R2R3-MYB domains. Beyond these domains, the amino acid sequences diverged greatly for different MYB308 homologs (Figure 1B). We fused the coding sequence of *PlMYB308* without stop codon to the 5′ terminus of green fluorescent protein (GFP) to investigate the subcellular localization of PlMYB308. The recombinant construct pCAMBIA1301-PlMYB308-GFP was injected in the tobacco leaves. In tobacco cells, the PlMYB308-GFP fusion protein had strong signals in the nucleus and membrane, indicating that PlMYB308 was localized in the cell nucleus and membrane (Figure 1C).

**Figure 1.**
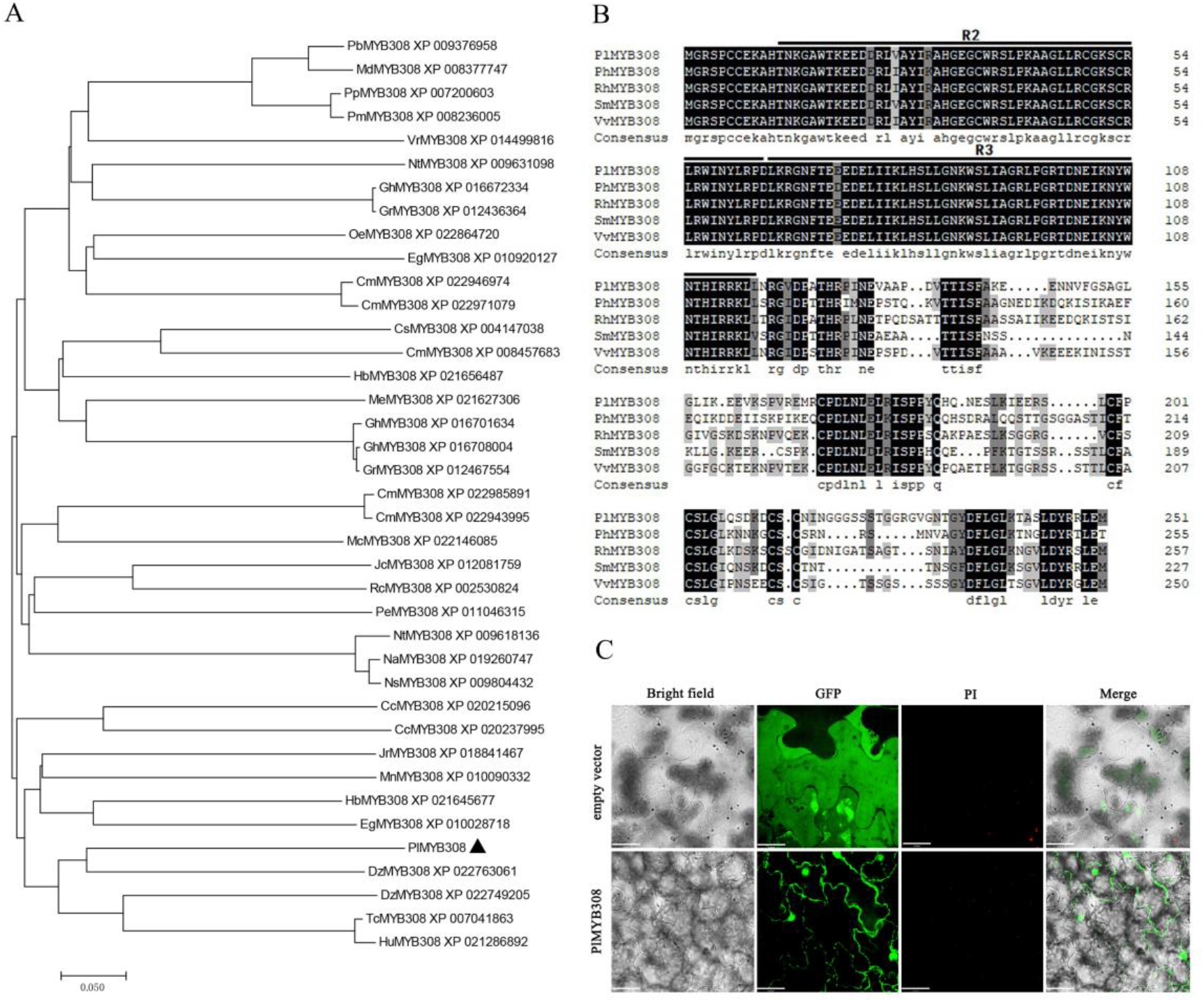
Phylogenetic analysis, multiple sequence alignment, and subcellular localization. (A) Phylogenetic tree generated using various MYB308 protein sequences from *Eucalyptus grandis, Ricinus communis, Fragaria vesca* L. and other plant species. Bootstrap values indicated the divergence of each branch, the scale indicated branch length. (B) Homologous sequences compared using various MYB308 protein sequences from *Paeonia lactiflora, Petunia hybrida, Salvia miltiorrhiza*, and *Vitis vinifera*. (C) Subcellular localization of GFP fusion of PlMYB308. Scale bars, 32 µm.

The complete flower senescence process of herbaceous peony was shown in Figure S1. The data of quantitative real-time PCR (qRT-PCR) showed that transcript abundance of *PlMYB308* significantly increased during flower senescence and reached the highest level at 6 days after anthesis (Figure 2A). This process was accompanied by the increase in transcript levels of senescence-related gene *PlSAG12* and the decrease in accumulation levels of anthocyanin (Figure 2, B and C). This implied that *PlMYB308* possibly plays an important role in flower senescence and anthocyanin synthesis.

**Figure 2.**
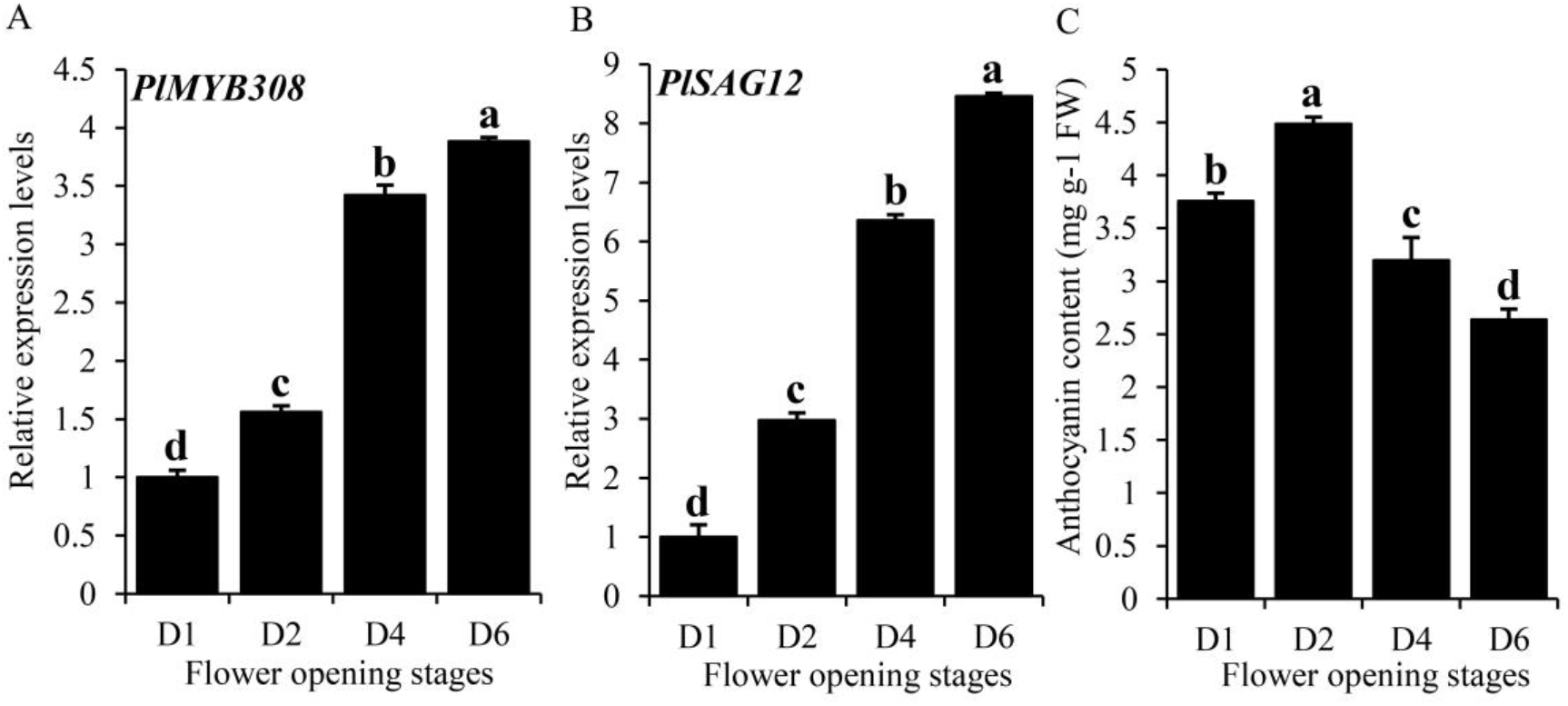
Expression levels of *PlMYB308* and *PlSAG12* and anthocyanin content during flower senescence. (A), (B) Quantitative real-time PCR analysis of *PlMYB308* and *PlSAG12* expression levels in herbaceous peony petals at D1, D2, D4, D6 after anthesis. *PlUB* was used as an internal control. (C) Anthocyanin content in herbaceous peony petals at D1, D2, D4, D6 after anthesis were determined. Error bars showed SD of the means of three biological replicates. Different letters indicate significant difference at P ≤ 0.05 by Duncan’s test.

### Silencing or overexpression of *PlMYB308* influences flower longevity

To uncover the function of *PlMYB308* in petal senescence, we used a virus-induced gene silencing (VIGS) approach to silence *PlMYB308* in herbaceous peony petal discs and overexpressed *PlMYB308* in tobacco plants using a stable genetic transformation technique. A 199-bp fragment of *PlMYB308* cDNA was introduced into the tobacco rattle virus (TRV) vector to generate the TRV-*PlMYB308* construct, which was used to silence *PlMYB308* in herbaceous peony petal discs. The phenotype of petal color fading started at 2 days (D2) after infiltration with empty vector, and the petal discs almost turned brown at 6 days (D6). By contrast, the *PlMYB308*-silenced discs only had slight color fading at D6 after inoculation (Figure 3A), exhibiting a clearly delayed senescence phenotype. Transcript abundance of *PlMYB308* remarkably reduced in TRV-*PlMYB308-*infiltrated petal discs, compared with TRV empty vector controls (Figure 3B). Approximately 3% of petal discs began to decay at D2, and the decay rate reached 66% at D6 in the controls. But in the *PlMYB308*-silenced discs, relatively lower petal decay rates were observed at D2 (1%) and D6 (32%) after inoculation (Figure 3C). Moreover, expression levels of *PlSAG12* in the *PlMYB308*-silenced petal discs were significantly lower than those in the TRV empty vector controls (Figure 3D).

**Figure 3.**
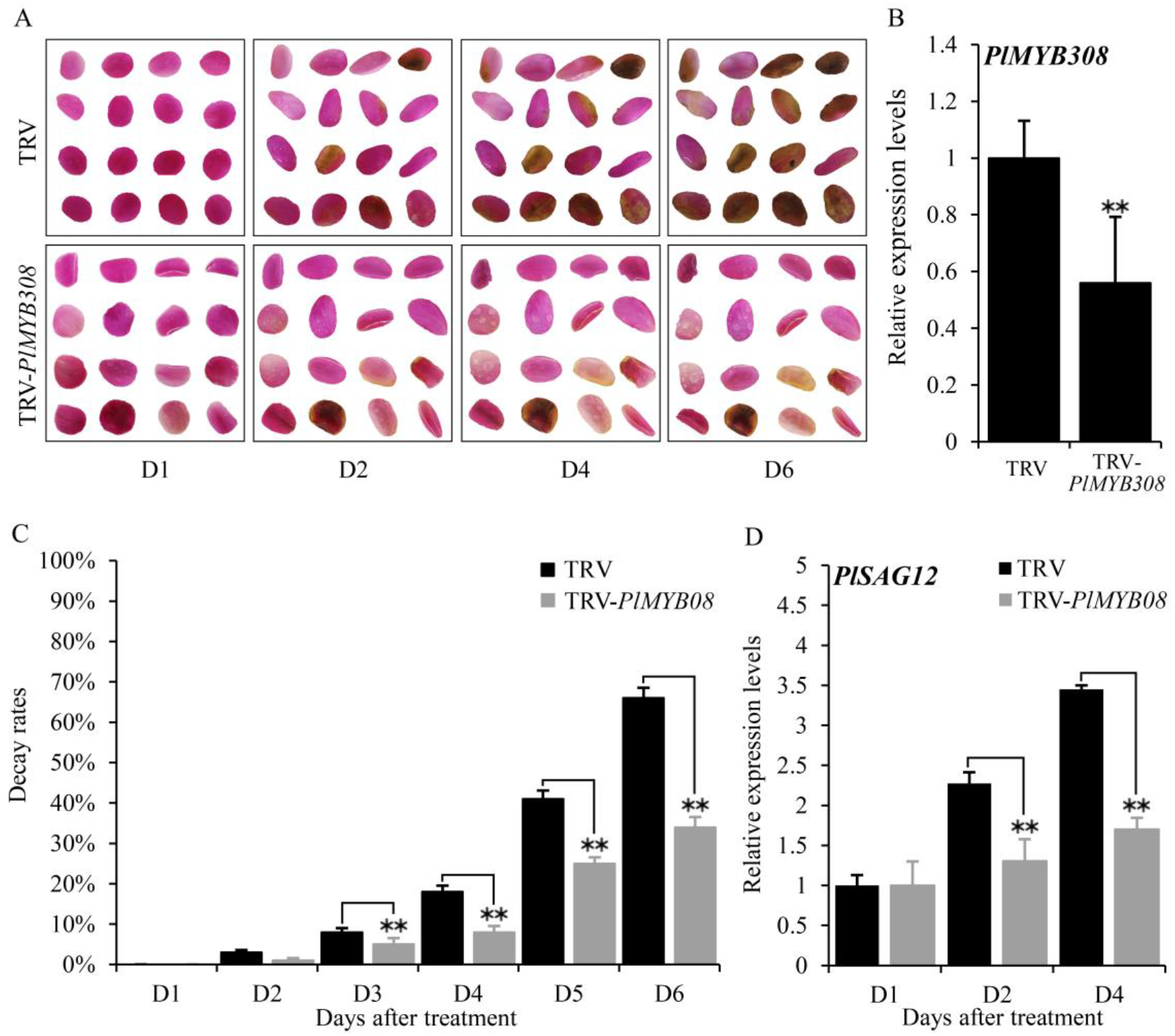
*PlMYB308* silencing delayed senescence of herbaceous peony petal discs. (A) The phenotypes of the petal discs were photographed at D1, D2, D4, D6 after infiltration with TRV empty vector and TRV-*PlMYB308*. (B) Abundances of *PlMYB308* transcript in empty vector- and TRV-*PlMYB308*-infected petal discs at D4. *PlUB* was used as an internal control. (C) Decay rates of the petal discs infiltrated with TRV empty vector and TRV-*PlMYB308*. (D) Expression levels of *PlSAG12* by quantitative real-time PCR in empty vector- and TRV-*PlMYB308-*infiltrated petal discs. Error bars showed SD of the means of three biological replicates. Asterisks indicated statistically significant differences by Student’s *t* test (\**P* < 0.05, \*\**P* < 0.01).

We transformed the model plant tobacco with a full-length coding sequence of *PlMYB308* cDNA. Transgenic tobacco plants displayed visibly shorter stems and smaller leaves than the wild-type (WT) plants (Figure 4, A and B). Different transgenic lines (#2, #5, and #7) showed accelerated flower senescence (Figure 4, C and E). Strong constitutive expression of *PlMYB308* was found in three transgenic lines (Figure 4D). Particularly, the line (#5) exhibiting the maximum expression of *PlMYB308* was selected for further analysis. In *PlMYB308-*overexpressing transgenic plants, the abundance of *NtSAG12* transcript was significantly higher than that in the WT plants during flower senescence (Figure 4F).

**Figure 4.**
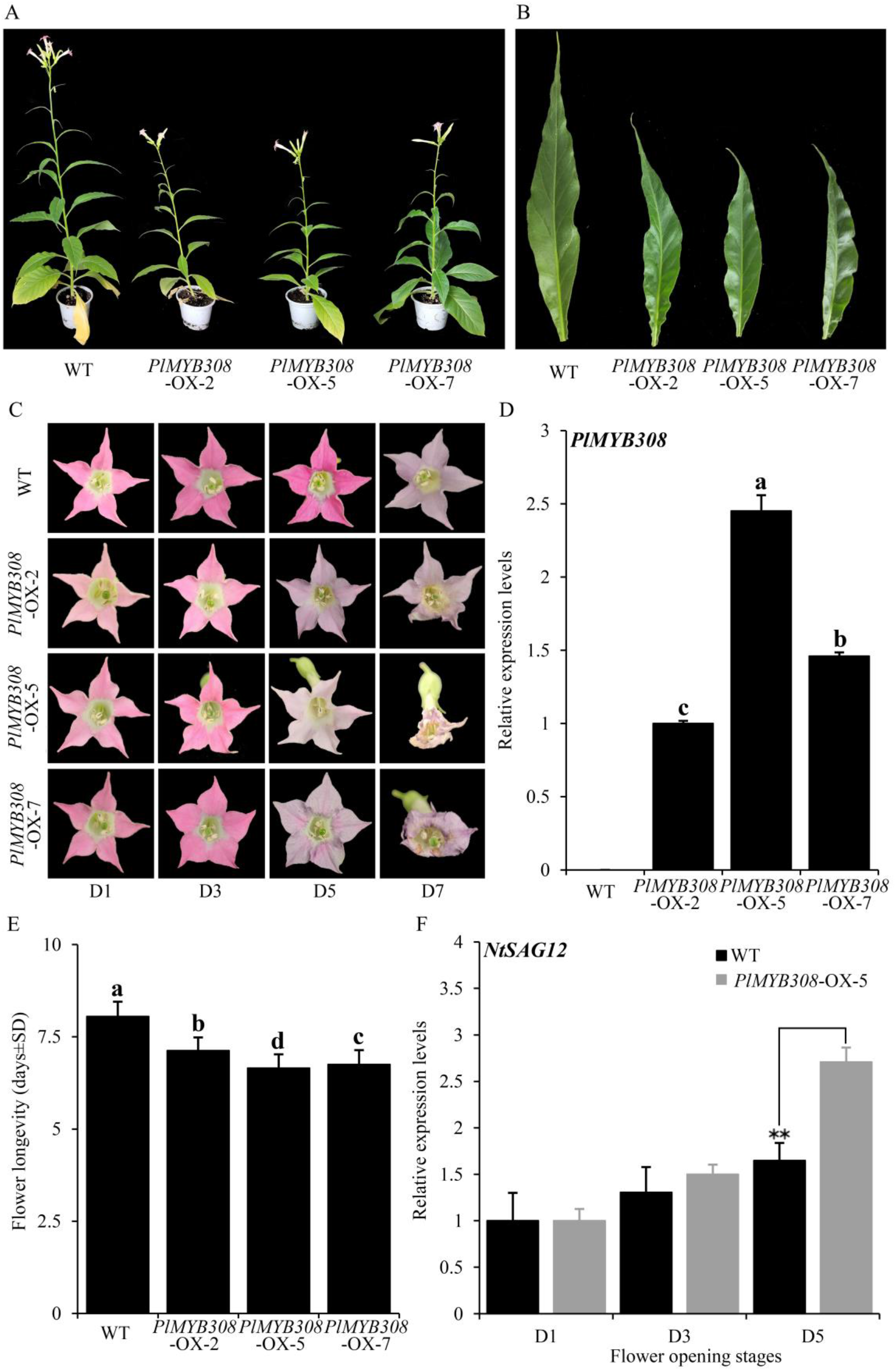
*PlMYB308* overexpression accelerated flower senescence in tobacco. (A)(B) *PlMYB308* overexpression led to shorter stems and smaller leaves than the wild-type (WT) control. Ten-week-old plants of WT and *PlMYB308*-overexpressing lines (OX-2, OX-5, and OX-7) were recorded and photographed. The fully expanded leaves were collected from the second leaves from the top of the branch. (C) The floral phenotypes were photographed at D1, D3, D5 and D7 after anthesis. (D) Expression levels of *PlMYB308* in wild-type (WT) plants and *PlMYB308-*overexpressing transgenic plants (*PlMYB308*-OX) by quantitative real-time PCR at D1. *PlMYB308*-OX-2, *PlMYB308*-OX-5 and *PlMYB308*-OX-7, different lines of *PlMYB308* overexpression. (E) Longevities of attached flowers in WT and *PlMYB308*-overexpressing transgenic tobacco plants. Means±SD for 60 flowers (15 from each of four replicate plants). (F) Expression levels of *NtSAG12* in WT and *PlMYB308*-overexpressing line (OX-5) at given time points. Error bars show SD of the means of three biological replicates. Asterisks indicated statistically significant differences (\**P* < 0.05, \*\**P* < 0.01, Student’s *t* test).

### *PlMYB308* affects ethylene, ABA, and GA production

As ethylene, ABA, and GA are important hormones for regulating flower senescence, we investigated their accumulation in *PlMYB308-*silenced herbaceous peony petal discs and *PlMYB308-*overexpressing transgenic tobacco plants. GA production was significantly higher in *PlMYB308*-silenced petal discs at 4 days (D4) after infiltration compared with the TRV empty vector controls (Figure 5A). On the contrary, GA levels were much lower in *PlMYB308*-overexpressing flowers at 5 days (D5) after anthesis than that in WT flowers (Figure 5D). Ethylene accumulation was markedly lower in petal discs with *PlMYB308* silencing but higher in transgenic tobacco lines with *PlMYB308* overexpression, compared to the controls (Figure 5, B and E). The content of ABA showed the same trends as that of ethylene in *PlMYB308*-silenced or - overexpressing plants (Figure 5, C and F). These results suggested that PlMYB308 regulated flower senescence by modulating the ethylene, ABA, and GA biosynthesis.

**Figure 5.**
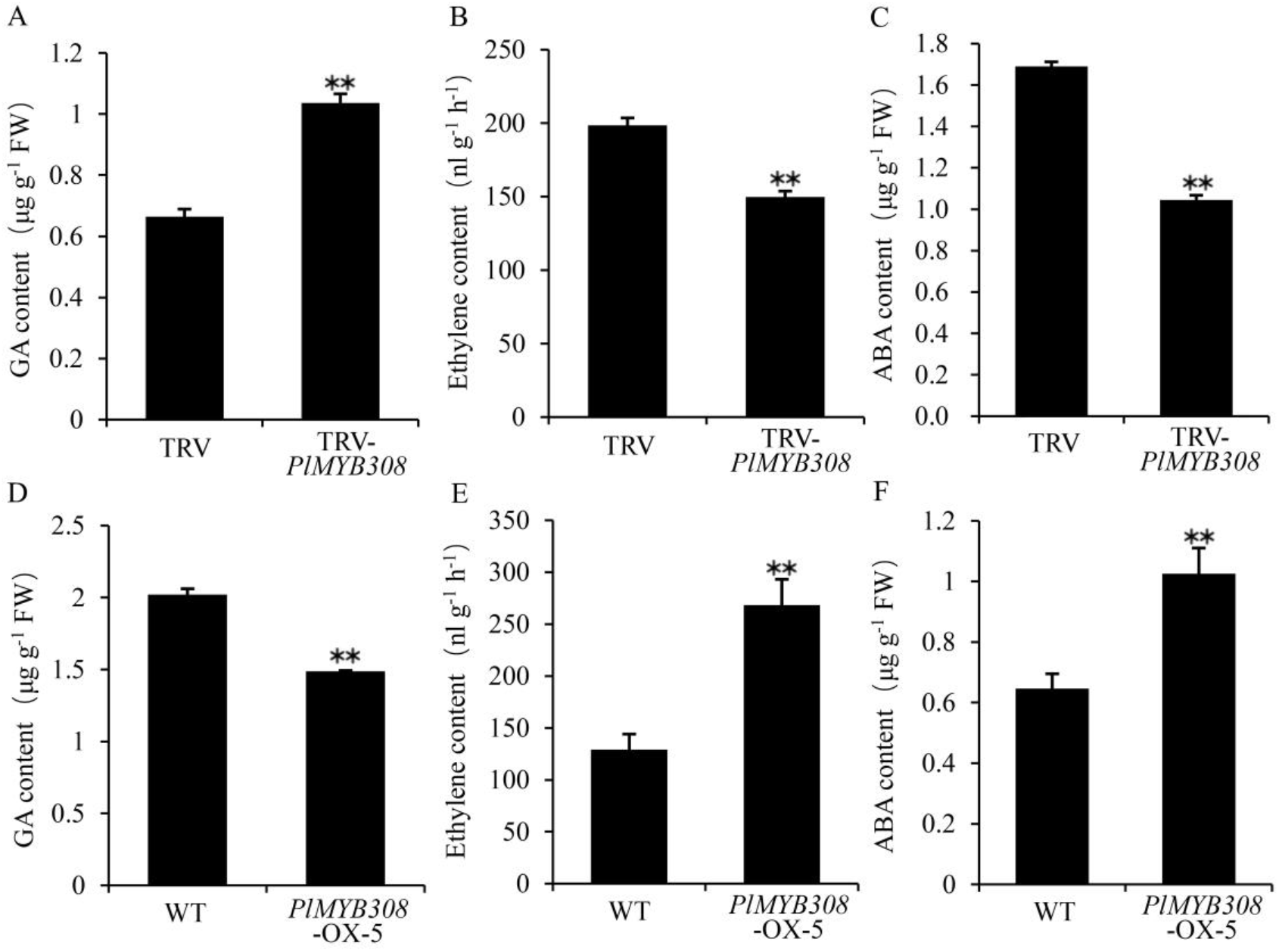
Effect of overexpression and silencing of *PlMYB308* on GA, ethylene and ABA production. (A), (B), (C) Content of GA, ethylene, and ABA in TRV empty vector- and TRV-*PlMYB308*-infiltrated herbaceous peony petal discs. GA, ethylene, and ABA production was measured at D4 after infiltration. (D), (E), (F) Content of GA, ethylene and ABA in wild-type and *PlMYB308*-overexpressing transgenic tobacco line (OX-5). GA, ethylene, and ABA production was measured at D5 after anthesis. Error bars show SD of the means of three biological replicates. Asterisks indicate statistically significant differences (\**P* < 0.05, \*\**P* < 0.01, Student’s *t* test).

### *PlMYB308* regulates the expression of some hormone biosynthetic genes

To further understand the role of *PlMYB308* in regulating hormone accumulation, we examined transcript levels of 12 different genes from herbaceous peony and tobacco plants related to hormone biosynthesis. Comparing to the controls, *PlGA20ox1* and *PlGA3ox1* were significantly up-regulated in *PlMYB308*-silenced petal discs while *PlGA2ox1, PlACO1, PlACO3, PlACS1, PlACS7*, and *PlNCED2* were down-regulated (Figure 6A). On the other hand, *NtGA3ox1* expression levels substantially decreased in *PlMYB308*-overexpressing transgenic tobacco lines compared to the WT, whereas transcript abundances of *NtACO1, NtACO3, NtACS1, NtAAO*, and *NtNCED2* increased (Figure 6B). These gene expression results were in agreement with that obtained from the hormone measurement above.

**Figure 6.**
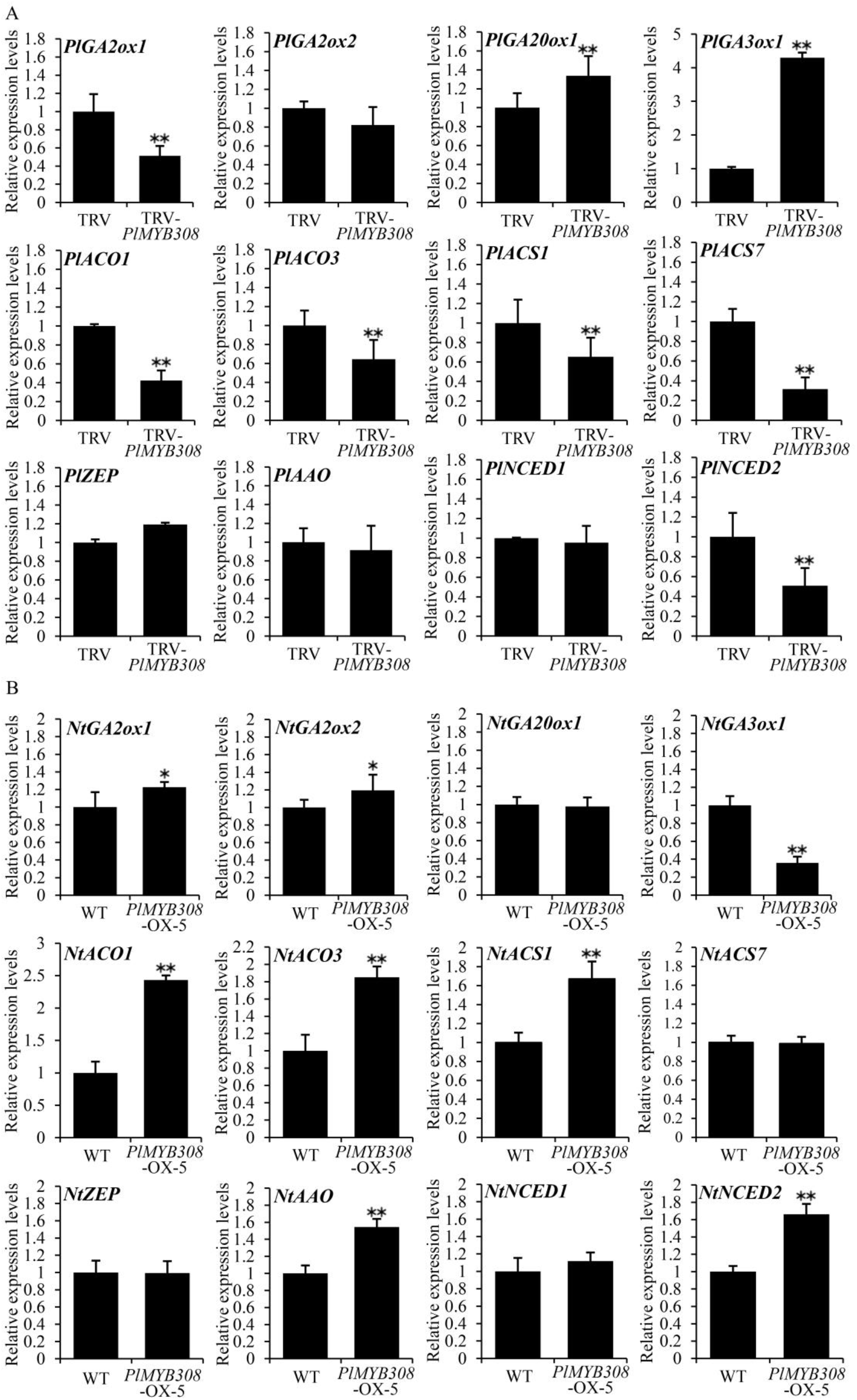
Expression analysis of hormone biosynthesis -related genes. (A) The petal discs were sampled for gene expression analysis at D4 after infiltration with TRV empty vector and TRV-*PlMYB308. PlUB* was used as an internal control. Relative expression levels were normalized to the empty vector control. (B) The tobacco flowers were sampled for gene expressions analysis at D5 after anthesis. *NtEF-1α* was used as an internal control. Relative expression levels were normalized to the WT. Error bars show SD of the means of three biological replicates. Asterisks indicate statistically significant differences (\**P* < 0.05, \*\**P* < 0.01, Student’s *t* test).

### PlMYB308 binds to the promoter of ethylene biosynthetic gene *PlACO1*

As we described above, the promoters of ethylene, ABA, and GA biosynthetic genes *PlGA3ox1, PlACO1, PlACO3, PlACS1*, and *PlNCED2* were isolated as the potential targets of PlMYB308 transcriptional activation. Their promoter sequences were submitted to the online website PlantCare for predictive analysis, revealing a diversity of *cis*-acting elements within the promoter regions (Table S1-5).

To determine whether the genes associated with ethylene, ABA and GA biosynthesis were targeted by PlMYB308, a dual-luciferase assay was subsequently carried out. The pGreenII 62-SK-PlMYB308 recombinant plasmid was constructed to co-infiltrate with the pGreenII 0800 constructs harboring different promoter sequences, and the control was the pGreenII 62-SK-empty vector (Figure 7A). The co-expression of pGreenII 62-SK-*PlMYB308* with pGreenII 0800-*PlACO1* promoter resulted in a 2.5-fold rise in firefly luciferase (LUC) activity, demonstrating that PlMYB308 significantly activated the promoter of *PlACO1* (Figure 7B). These data suggested that PlMYB308 may be a specific regulator of ethylene biosynthesis by targeting *PlACO1*.

**Figure 7.**
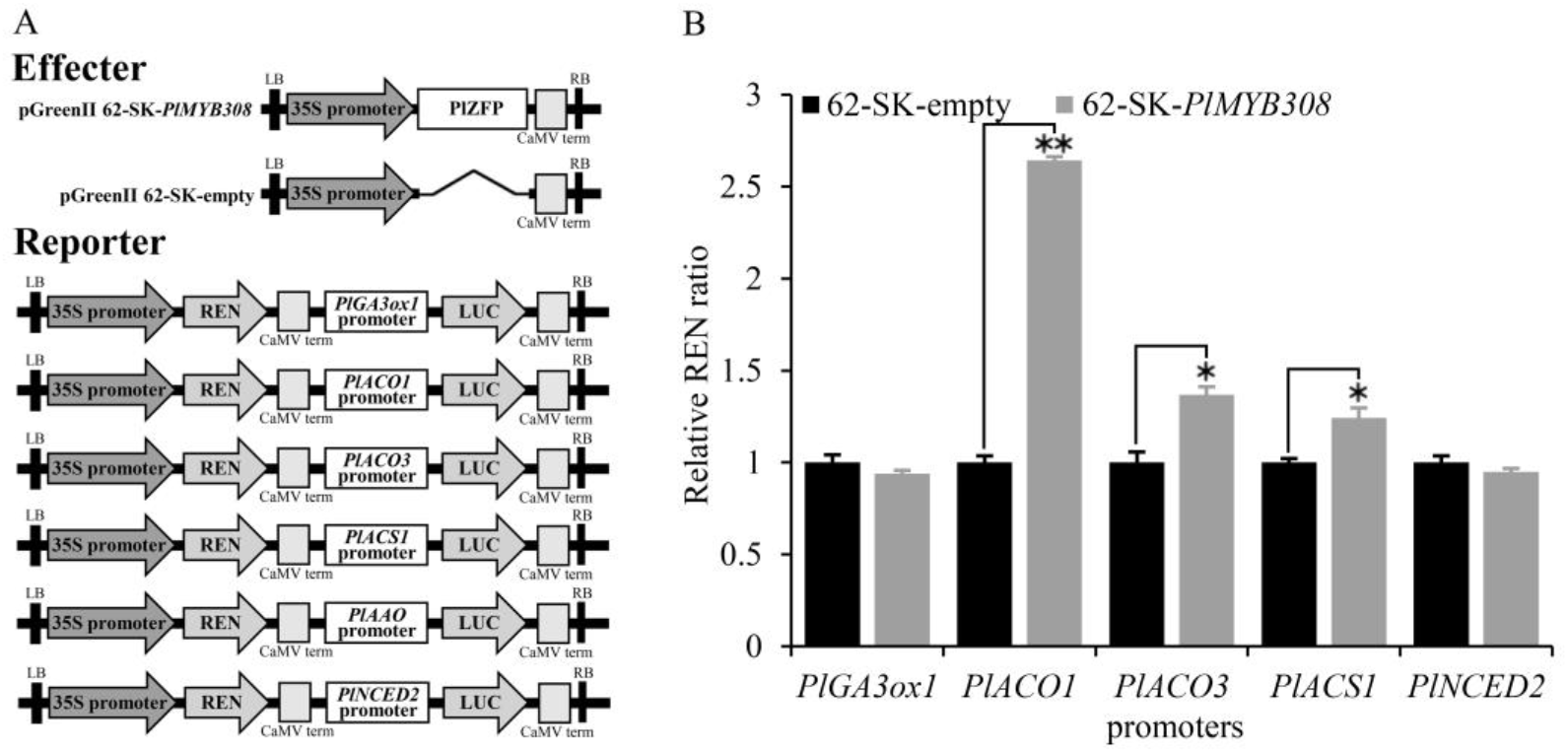
Transient dual-luciferase reporter assay. (A) Schematic diagrams of the reporter and effector constructs used for the dual-luciferase assay. (B) PlMYB308 transactivated the promoter of *PlACO1*. The ratio of LUC/REN of the empty vector (62-SK) plus promoter was considered as a calibrator (set as 1). The activation is indicated by the ratio of LUC to REN. Data represent the mean ± SD of six independent repeats. ** indicates statistically significant differences at the level of *P* < 0.01 by Student’s *t* test.

### PlMYB308 interacts with putative anthocyanin negative regulator PlbHLH33

In this study, the flower senescence and color-fading phenotypes in herbaceous peony or tobacco always co-existed. Next, we examined the correlation of anthocyanin content with *PlMYB308* expression levels in *PlMYB308*-silenced herbaceous peony petal discs and *PlMYB308*-overexpressing transgenic tobacco flowers (Figure 8, A– D). The anthocyanin content decreased with the increasing expression levels of *PlMYB308* at various senescing phases, which represented a negative correlation between anthocyanin accumulation and *PlMYB308* transcription. Amino acid analysis showed that PlMYB308 possesses a conserved bHLH binding site (Figure S3). It can be inferred that there may be an interaction between PlMYB308 and certain PlbHLHs. It is known that bHLH33 regulates anthocyanin biosynthesis in some plant species (Liu et al. 2019). Our further bimolecular fluorescence complementation experiment showed a remarkable interaction of PlMYB308 with PlbHLH33 in the cell nucleus (Figure 8E). The interaction between them may impose a negative influence on anthocyanin biosynthesis.

**Figure 8.**
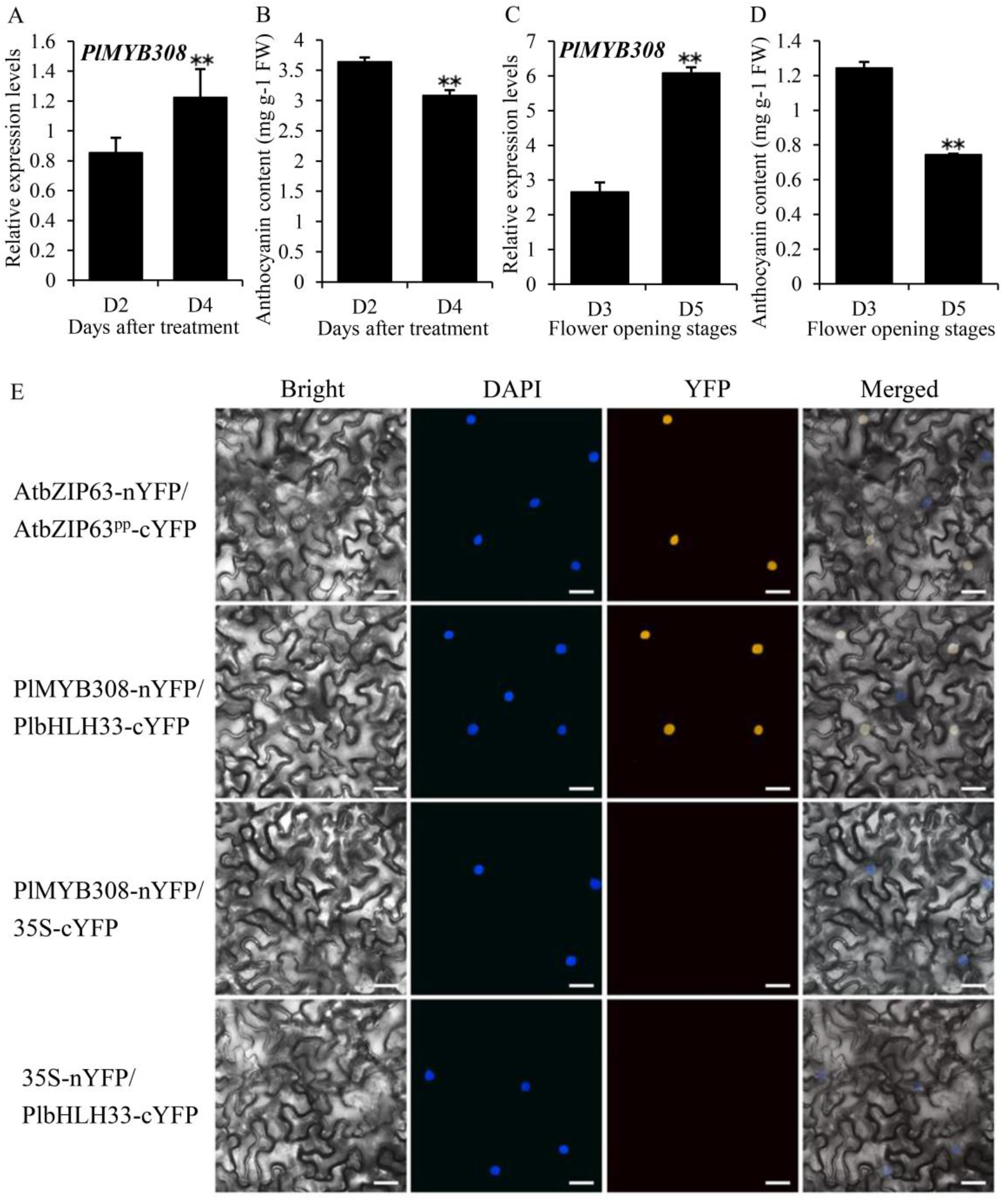
Interaction of PlMYB308 with PlbHLH33. (A) *PlMYB308* expression levels in *PlMYB308*-silenced herbaceous peony petal discs at D2 and D4 after treatment. (B) Anthocyanin content in *PlMYB308*-Silenced herbaceous peony petal discs at D2 and D4 after treatment. (C) *PlMYB308* expression levels in *PlMYB308*-overexpressing transgenic tobacco flowers at D3 and D5 after anthesis. (D) Anthocyanin content in *PlMYB308*-overexpressing transgenic tobacco flowers at D3 and D5 after anthesis. (E) Bimolecular fluorescence complementation assay of the interaction between PlMYB308 and PlBHLH33 in tobacco cells. Scale bars, 25 µm. AtbZIP63 and AtbZIP63^pp^ were used as a positive control (Walter et al. 2004).

## Discussion

Flower senescence is a complex biological process and always concurrent with the phenomenon of pigment fading. It is well established that plant hormones normally work in combination to play important roles in flower senescence and anthocyanin accumulation (Zhang and Zhou, 2013). We identified a MYB TF namely PlMYB308 in herbaceous peony in this study, and its amino acid sequence shows high similarity to MYB308 proteins in other plant species. PlMYB308 regulated herbaceous peony petal senescence by inducing ethylene and ABA but inhibiting GA production. The data presented here demonstrated that PlMYB308 bound the promoter of an ethylene biosynthetic gene, *PlACO1*, and thereby specifically activated the expression levels of *PlACO1*. We also found that PlMYB308 could interact with PlbHLH33, which probably negatively regulated the flower pigmentation.

### PlMYB308 is involved in flower senescence by activating *PlACO1* expression in herbaceous peony

The MYB protein family is composed of a massive number of members in all eukaryotes. Here, we showed that *PlMYB308*, a R2R3 MYB TF from herbaceous peony, was significantly up-regulated in different periods of flower senescence (Figure 2A). Because the stable genetic transformation system of herbaceous peony has not been established, we used tobacco as a heterologous expression model system for studying the function of PlMYB308. Overexpression and silencing of *PlMYB308* reduced and increased flower longevity, respectively, suggesting that PlMYB308 may be an essential regulator of flower senescence. It is well recognized that plant hormones play synergistic or antagonistic roles in the modulation of flower senescence (Zhang and Zhou, 2013). In many ornamental crops, such as *Lathyrus odoratus, Gypsophila paniculata*, and *Antirrhinum majus*, a climacteric rise in ethylene production initiates flower senescence (Woltering et al., 1996). ABA imposes an analogous influence with ethylene on flower senescence. In carnation, the content of ABA begins to increase after harvest of cut-flowers, and exogenous ABA also enhances the accumulation of endogenous ABA and accelerates the flower senescence (Onoue et al., 2000). Nevertheless, GA has an opposite effect on senescing process of floral tissues compared to ethylene and ABA. In rose flowers, the treatment of GA_3_ could maintain the integrity of cell membrane, thus delaying the occurrence of senescence (Saks and van Staden, 1993). Our research showed that overexpression and silencing of *PlMYB308* influenced the production of ethylene, ABA, and GA in petals, suggesting that PlMYB308 positively regulates flower senescence by altering endogenous hormones content.

The biosynthesis pathways of various hormones have been extensively studied in plants. In the GA biosynthesis pathway, GA20-oxidase and GA3-oxidase are two important enzymes catalyzing the steps of bioactive GA production (Spray et al., 1996). Zeaxanthin epoxidase (ZEP), 9-cis-epoxycarotenoid dioxygenase (NCED), and aldehyde oxidase (AAO) are indispensible participants of ABA biosynthesis in higher plants (Zhang DP, 2014). 1-aminocyclopropane-1-carboxylate synthase (ACS) and ACC oxidase (ACO) are two key enzymes required for ethylene synthesis (Zarembinski et al., 1994). We detected the transcript abundances of 12 hormone biosynthetic genes, and found that transcript levels of *GA3ox1, ACO1, ACO3, ACS1*, and *NCED2* were variable in *PlMYB308*-silenced or -overexpressing plants. Furthermore, the dual-luciferase assay revealed a direct binding of PlMYB308 to the promoter of *PlACO1* (Figure 7B). Our study demonstrates that PlMYB308 may regulate flower senescence by targeting the ethylene biosynthesis pathway.

The change in transcript levels is one of the most important regulatory modes in organisms. The predictive analysis of *PlACO1* promoter sequence showed 5 *cis*-acting elements, including Box4, G-Box, GT1-motif, MYB, and TCT-motif. Among them, G-Box is a 6-bp CACGAC *cis*-acting element that exists widely in the promoter regions responsive to ABA, jasmonic acid, ethylene, elicifor, and hypoxia signals (Shen et al., 1996). In addition, tobacco MYB305 was shown by an electrophoretic mobility shift assay to bind to the G-Box element in the *NtACO1* promoter (Sablowski et al., 1995). This report showed that MYB proteins specifically bind to the G-Box motifs, which is in accordance with our findings that PlMYB308 directly targeted the G-Box-containing promoter of *PlACO1*.

### PlMYB308 may be participated in the crosstalk among ethylene, ABA, and GA during flower senescence

Ethylene, ABA, and GA are all involved in plant growth and development processes such as stomata formation, seed germination, and flower senescence (Richards et al., 2001; Fleet et al., 2005). The crosstalk between these three hormones has been demonstrated in many plants. Active ethylene signaling results in decreased GA production. Previous evidence suggests that the reduction of GA levels enhances ethylene-mediated flower senescence in rose (Lü et al., 2014). ABA and GA have obvious antagonistic actions in regulating plant growth and development. In tobacco, ABA and GA jointly contribute to the leaf senescence process. When GA accumulation is kept at a high level, tobacco leaves will not quickly enter the aging state (Wei et al., 2020). Both ethylene and ABA play important roles in the senescence of cut flowers, and they can promote the synthesis of each other (Doorn and Woltering, 2008). In this study, in herbaceous peony, GA_3_ treatment remarkably delayed petal senescence which was induced by ABA and ethylene (Figure S2). In *PlMYB308*-overexpressing transgenic tobacco plants, ethylene and ABA production was higher and GA production was lower than the WT plants (Figure 5, D–F). As we discussed before, PlMYB308 is involved in herbaceous peony flower senescence by promoting *PlACO1* expression in herbaceous peony. We supposed that overexpression of *PlMYB308* initially accelerated ethylene biosynthesis, followed by a promotion of ABA accumulation. Thus, the two hormones collaboratively resulted in short longevity of flowers (Figure 4, C and E). On the other hand, the enhanced ethylene release inhibited GA production, and thereby leading to the repressed growth phenotypes in transgenic tobacco plants (Figure 4, A and B). By contrast, when silencing *PlMYB308* in herbaceous peony petals, ethylene and ABA production decreased while GA levels increased (Figure 5, A–C). This could be explained that the impaired ethylene production by down-regulation of *PlMYB308* interrupted the ABA biosynthesis but facilitated GA production. These data demonstrate that PlMYB308 mediates the relationship of ethylene, ABA, and GA in the positive or negative manner. With the exception of *PlACO1*, however, we found no exact structural genes associated with ABA or GA biosynthesis binding to PlMYB308 in our work. It would be quite necessary to comprehensively analyze the ABA and GA biosynthetic genes in *PlMYB308*-silenced and -overexpressing plants in the future.

### PlMYB308 simultaneously acts on ethylene and anthocyanin accumulation and ethylene may inhibit anthocyanin biosynthesis in herbaceous peony

The senescence with suppressed pigmentation is common in herbaceous peony flowers. It is noteworthy that the increase in ethylene production can boost senescence and trigger color fading. In some fruits, such as grape (Kereamy et al., 2003), ethylene promotes anthocyanin biosynthesis. However, some present investigations revealed that in Arabidopsis, ethylene can be an inhibitor of anthocyanin biosynthesis (Jeong et al., 2010). The molecular mechanism underlying ethylene biosynthesis and anthocyanin accumulation remains enigmatic in flower senescence. Some R2R3-MYB TFs which regulate anthocyanin accumulation always interact with bHLH TFs.

In our study, PlMYB308 can interact with PlbHLH33 (Figure 8E), whose homologs in other species have been described as a negative regulator of anthocyanin production (Liu et al. 2019). This suggests that PlMYB308 may inhibit anthocyanin biosynthesis in herbaceous peony via a co-operation with a unique bHLH TF. As we mentioned above, PlMYB308 regulated ethylene biosynthesis by binding to the promoter of *PlACO1*, and the pigment production by interacting with PlbHLH33. Accordingly, we concluded that ethylene perhaps inhibits anthocyanin biosynthesis through the transcriptional regulation of PlMYB308 in herbaceous peony flowers. Ultimately, a model was proposed that PlMYB308 acts on flower senescence of herbaceous peony by modulating anthocyanin accumulation and hormone crosstalk among ethylene, ABA, and GA (Figure 9). This model can show why flower senescence of herbaceous peony co-exists with pigment fading.

**Figure 9.**
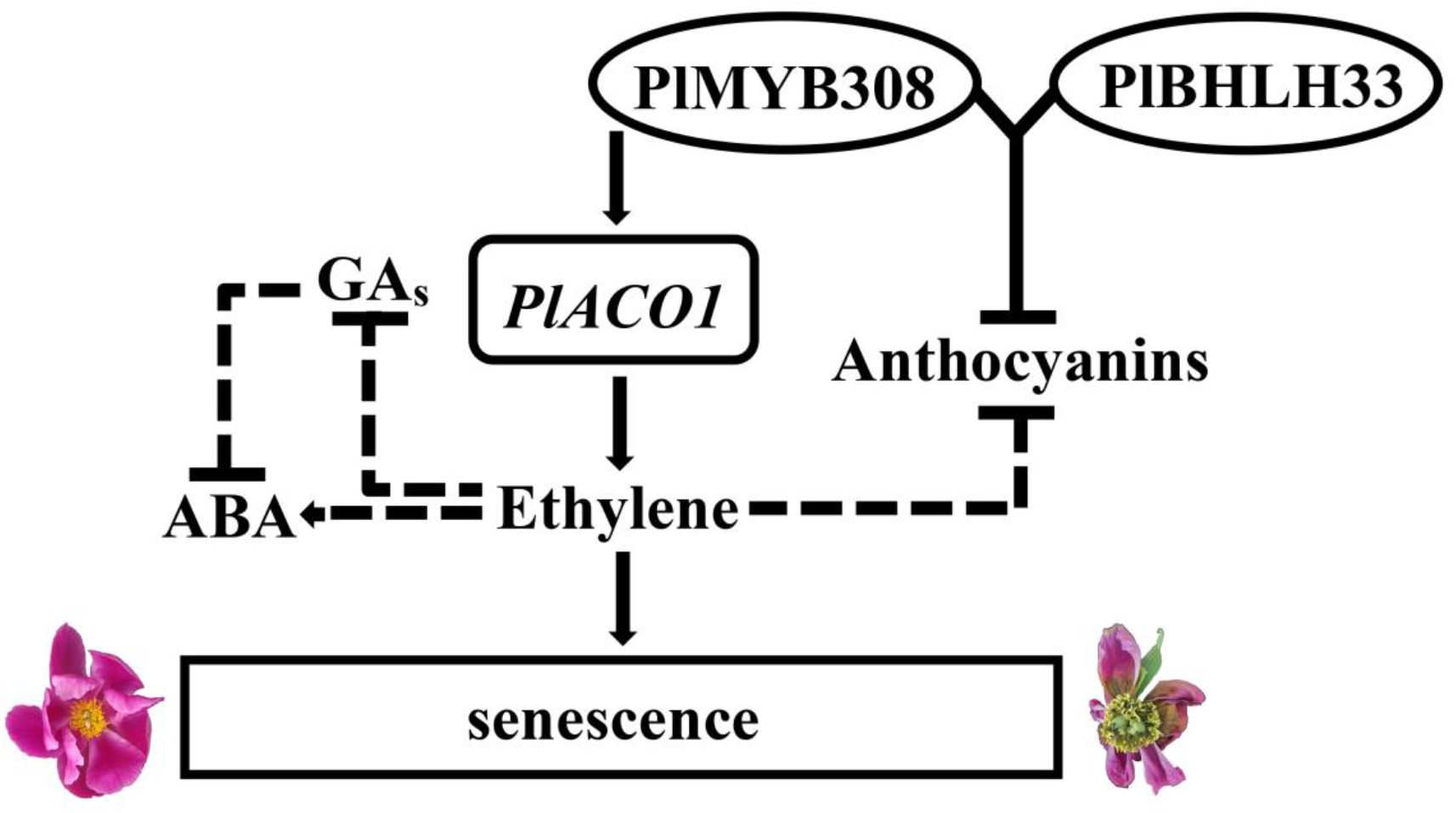
A proposed model of PlMYB308 regulating flower senescence and anthocyanin accumulation. Arrows indicate promotion, and leading dashes indicate inhibition.

## Materials and methods

### Plant materials and growth conditions

To avoid the variation in gene expression levels in multi-layered petals, a herbaceous peony cultivar (*Paeonia lactiflora* ‘hangshao’) with single-layered petals was used in this study. Plants were grown in the germplasm resource garden of Northwest Agriculture and Forestry University. The clean floral samples were collected at the first day after flowering. The samples were kept in deionized water for the VIGS assay to silence *PlMYB308* (Ma et al., 2005). The petals used for expression assessment were collected at four developmental stages: the first day after flowering (D1); the second day after flowering (D2); the fourth day after flowering (D4); the sixth day after flowering (D6). All samples were immediately frozen in liquid nitrogen and stored at -80ºC (Du et al., 2015). The seeds of tobacco (*Nicotiana tabacum*) were germinated at 24/20°C day/night temperature, 16/8h light/dark cycle and a 70% relative humidity. These plants were used for ectopic overexpression experiment of *PlMYB308*.

### Identification of *PlMYB308*

We use TIANGEN RNA Prep Pure Plant kit (Tiangen, Beijing, China) to isolated the total RNA of herbaceous peony petals. The first-strand cDNA was synthesized using a PrimeScript® RT reagent Kit (Takara, Otsu, Shiga, Japan). The *PlMYB308* gene was PCR-amplified using specific primers (Table S6). The amplified gene sequence was verified by TSINGKE. We use DNAMAN software (version 8.0) to compare the homologous sequence of PlMYB308 in herbaceous peony and other species and use Neighbor-Joining method (NJ) of MEGA software (version 7.0) to construct the phylogenetic tree. The conserved domain was identified by SMART (http://smart.embl-heidelberg.de/smart/set_mode.cgi?NORMAL=1).

### Subcellular localization assay

To determine where the protein was expressed in the cells, we constructed a *PlMYB308*-GFP fusion using the binary vector pCAMBIA1301-GFP. A biolistic PDS-1000He instrument (Bio-Rad, CA, USA) was used to bombard tobacco cells with this constructed plasmid. After incubation at 25°C for 16 h in the dark, the samples were observed for fluorescence signals under a confocal laser scanning microscope.

### Virus-induced gene silencing assay

Silencing of *PlMYB308* by TRV-based VIGS was performed as previously described by Dai et al. (2012). The TRV1, TRV2, and TRV2-*PlMYB308* constructs were transformed into *Agrobacterium tumefaciens* strain GV3101 individually, and then cultured them overnight. The culture mixture containing an equal ratio (v/v) of TRV1 and TRV empty vector as well as TRV1 and TRV-*PlMYB308* were used for the empty vector control and *PlMYB308* silencing experiments, respectively. Before vacuum infiltration, the mixture was shaken slightly at 28°C for 3 h. Herbaceous peony petals at D1 after anthesis were collected, and trimmed them into 0.8 mm discs in diameter from the center of the petals. The petal discs were immersed in the bacterial suspensions then infiltrated under a vacuum at 0.8 MPa for 15 min. Releasing vacuum and washing petal discs in deionized water then kept them for 24 h at 8°C. Kept these petal discs at 23°C and observed the phenotypes of them every day. Samples for RNA isolation were collected at the first day (D1); the second day (D2); the fourth day (D4); the sixth day (D6) after treatment.

### Plasmid construction and stable transformation of tobacco

To generate transgenic tobacco plants with *PlMYB308* overexpression, the complete ORF region of *PlMYB308* (a 759-bp cDNA sequence) was cloned into the pCAMBIA1300 vector between *Kpn*I and *Sal*I sites. *A. tumefaciens* strain LBA4404 was used to transformed the construct. PCR amplification and sequencing were conducted to confirm the integration of destination vector with the *PlMYB308*. Transformation and regeneration of tobacco plants were performed using the leaf disk method as previously described by Hong et al. (2013). In this study, we generated 9 independent transgenic lines at T1. Homozygous lines from T2 generation were used for the follow-up experiments.

### Quantitative real-time PCR assay

Quantitative real-time PCR (qRT-PCR) assay was used to determine the expression levels of *PlMYB308* and other flower senescence-associated genes in herbaceous peony and tobacco plants. The sequences of related genes were obtained from the NCBI Genbank database (https://www.ncbi.nlm.nih.gov/gene). We designed specific primers according to the cDNA sequences of all genes tested (Table S6). SYBR Premix Ex Taq II (Takara, Otsu, Shiga, Japan) on StepOnePlus Real Time PCR system (Thermo Fisher Scientific, Waltham, MA, USA) was used for the qRT-PCR assay. We used *PlUB* and *NtEF-1α* as internal controls to normalize the expression data in herbaceous peony and tobacco respectively. At the end of qRT-PCR program, the melting curve program was used to ensure the specific amplification. We used the 2^-ΔΔCT^ comparative threshold cycle (Ct) method to calculate the relative expression levels of genes. Three biological replicates were used for each measurement.

### Decay rates and flower longevity

Decay rates of the herbaceous peony petal discs were recorded per day after inoculation. We recorded flower longevity from anthesis to complete wilting. Collected one hundred petal discs from TRV empty vector control petal discs (TRV) and in *PlMYB308* silenced petal discs (TRV-*PlMYB308*) respectively from 1 day to 6 days to detect the decay rates of herbaceous peony petal discs in VIGS assay. 15 flowers from wild-type (WT) plants and *PlMYB308-*OX-2, *PlMYB308-*OX-5, *PlMYB308-*OX-7 transgenic plants respectively were used for the detection of decay rates and flower longevity.

### Anthocyanin content measurement

Anthocyanin levels in herbaceous peony petal discs and tobacco corollas were determined according to Luo et al (2017) method. Added liquid nitrogen into about 100 mg of samples and grounded them into fine powders. Used 1% HCl-methanol solution to extract the anthocyanin at 4°C for 24 h and then ultrasound for 60 min. 12,000 rpm centrifugated for 10 min and then purified the supernatant through a 0.22 µm membrane filter. A spectrophotometer was used for absorbance determination at 530 nm and 657 nm. Used the equation ((A530 − 0.25 × A657) × FW^−1^) to calculated anthocyanin content. Three biological replicates were used for each measurement.

### Measurement of ethylene production

Ethylene production was measured at D6 after infiltration with TRV empty vector control and TRV-*PlMYB308* in petal discs of herbaceous peony, and at D5 in WT and *PlMYB308*-overexpressing tobacco plants. About 100 mg of samples were placed into airtight tubes for 4 h at 26°C. Approximately 1 ml of gaseous samples were collected using a gastight hypodermic syringe and injected into a gas chromatographer (GC-8A; Shimadzu, Kyoto, Japan) for ethylene detection. Three biological replicates were analyzed for each sample.

### Measurement of GA and ABA production

GA and ABA production was measured as previously described by Pan et al. (2010). Added liquid nitrogen into about 100 mg of TRV empty vector- and TRV-*PlMYB308*-infected peony petal discs, and of WT and *PlMYB308*-overexpressing flowers samples respectively and grounded them into fine powders. Using extraction solvent (2:1:0.002 v/v/v 2–propanol/H_2_O/concentrated HCl; ratio of sample to solvent 1:10) extracted samples on a rotary shaker (100 rpm) at 4°C for 30 min and then added 1 ml of dichloromethane to each sample and shaken at 4°C for 30 min. 13 000 g centrifugated for 5 min at 4°C and then collected about 1.5 ml of samples at lower phase. Desiccated the solvent mixtures then re-dissolved them in 0.1 ml of methanol. The sample solution was detected by HPLC electrospray ionization tandem mass spectrometry (HPLC-ESI-MS/MS).

### Dual luciferase transient transfection assay

The pGreenII62-SK construct including the coding region of *PlMYB308*, and the pGreenII0800 constructs containing different promoter sequences were prepared as previously described by Zong et al. (2016) (Figure 7A). They function as the effector and reporter, respectively. Particularly, the reporter constructs harbored the gene promoters driving a *firefly luciferase* (*LUC*) gene and the CaMV 35 promoter driving a *Renilla luciferase* (*REN*) gene. Tobacco leaves were infected by the effector, reporter, and internal plasmid. Relative luciferase activities were assayed with the Dual-Luciferase Reporter® Assay System (Promega, Madison, WI, USA) using Promega GloMax 20/20 Microplate luminometer (Promega, Madison, WI, USA).

### Bimolecular fluorescence complementation assay

The PlMYB308-CYNP recombinant construct was produced by inserting the *PlMYB308* coding sequence into the pSPYNE-YFP vector. Meanwhile, the PlbHLH33-CYCP recombinant construct was generated by introducing the corresponding coding sequence into the pSPYCE-YFP vector. Transformed these two recombinant constructs into *A. tumefaciens* strain LBA4404. AtbZIP63 and AtbZIP63^pp^ were used as a positive control (Walter et al. 2004). Injected the *Agrobacterium* cultures to the Lower epidermis of tobacco leaves, and then incubated these tobaccos at 28°C for 3 days, and then observed under the LSM 510 Meta laser scanning confocal microscope (Zeiss).

## Supplemental data

**Supplemental Figure S1**. Different stages of herbaceous peony flower senescence.

**Supplemental Figure S2**. Senescence process of herbaceous peony petal disks treated with various hormones. After 24 h treatment with distilled water, 50 μM GA_3_, 200 μM ethylene, 100 μM ABA, combinations of 50 μM GA_3_ and 200 μM ethylene and combinations of 50 μM GA_3_ and 100 μM ABA respectively, the petal discs were kept on the wet filter papers and the photos were taken at first day (D1), second day (D2), fourth day (D4) and sixth day (D6) after treatment, respectively.

**Supplemental Figure S3**. bHLH binding site within the amino acids of PlMYB308.

**Supplemental Table S1**. *Cis*-acting component analysis in the promoter of *PlGA3ox1*.

**Supplemental Table S2**. *Cis*-acting component analysis in the promoter of *PlACO1*.

**Supplemental Table S3**. *Cis*-acting component analysis in the promoter of *PlACO3*.

**Supplemental Table S4**. *Cis*-acting component analysis in the promoter of *PlACS1*.

**Supplemental Table S5**. *Cis*-acting component analysis in the promoter of *PlNCED2*.

**Supplemental Table S6**. Genes specific primers used for gene isolation and expression analysis.

## Acknowledgments

We are grateful to Houhua Li for kindly providing the pSPYNE-YFP and pSPYCE-YFP vectors. We thank Jiaxin Meng for experimental assistance in dual luciferase assay. We are appreciative of Xiaokun Liu’s advice on operation of laboratory instruments.

